## Supplementary file (tables) for "Multimodal neural correlates of childhood psychopathology"

**Supplementary File 1**

Supplementary file 1a. Pearson’s correlations between PLS loadings (rows) obtained in the discovery sample and Michelini’s factor loadings (columns; Michelini et al. 2019). Significant correlations that survived FDR correction (*q* < 0.05) are indicated in bold. *Abbreviations: Intern = Internalizing, Extern = Externalizing, Neurodev = Neurodevelopmental.*

|  |  | **Michelini factors** | | | | | |
| --- | --- | --- | --- | --- | --- | --- | --- |
|  |  | P-factor | Extern | Intern | Neurodev | Somatoform | Detachment |
| LC1 | P-factor | **0.64** | **0.44** | **0.28** | 0.11 | **-0.39** | **-0.20** |
| LC2 | Intern / Extern | **-0.29** | **-0.80** | **0.62** | 0.04 | **0.32** | **0.26** |
| LC3 | Neurodev | -0.08 | **-0.17** | **-0.45** | **0.85** | **-0.26** | -0.04 |
| LC4 |  | -0.09 | -0.03 | **-0.30** | 0.14 | 0.11 | **0.30** |
| LC5 | Detachment | -0.02 | **-0.40** | 0.04 | **0.25** | -0.16 | **0.51** |

Supplementary file 1b. PLS loadings and z-scores for CBCL items for LCs 1-3. Significant loadings and z-scores that survived FDR correction (*q* < 0.05) are indicated in bold.

| **CBCL item** | **LC1 loading (z-score)** | **LC2 loading (z-score)** | **LC3 loading (z-score)** |
| --- | --- | --- | --- |
| Acts too young for his/her age | **0.46 (22.82)** | 0.03 (0.53) | **0.24 (3.31)** |
| Drinks alcohol without parents' approval | 0.07 (1.39) | -0.01 (-0.32) | **-0.09 (-2.13)** |
| Argues a lot | **0.58 (40.26)** | -0.08 (-1.21) | -0.11 (-1.47) |
| Fails to finish things he/she starts | **0.57 (38.81)** | 0.10 (1.40) | **0.24 (2.93)** |
| There is very little he/she enjoys | **0.45 (18.51)** | 0.09 (1.29) | -0.11 (-1.57) |
| Bowel movements outside toilet | **0.10 (3.79)** | -0.01 (-0.26) | 0.03 (0.81) |
| Bragging, boasting | **0.39 (18.10)** | **-0.10 (-1.81**) | -0.01 (-0.25) |
| Can't concentrate, can't pay attention for long | **0.59 (39.81)** | 0.09 (1.25) | **0.40 (4.70)** |
| Can't get his/her mind off certain thoughts; obsessions | **0.58 (35.14)** | **0.24 (3.28)** | 0.01 (0.18) |
| Can't sit still, restless, or hyperactive | **0.56 (33.03)** | 0.02 (0.31) | **0.26 (3.20)** |
| Clings to adults or too dependent | **0.45 (20.72)** | **0.21 (3.44)** | -0.05 (-0.74) |
| Complains of loneliness | **0.46 (20.38)** | **0.24 (3.84)** | **-0.13 (-1.92)** |
| Confused or seems to be in a fog | **0.42 (15.74)** | **0.21 (3.32)** | **0.18 (2.71)** |
| Cries a lot | **0.43 (19.13)** | **0.13 (2.02)** | **-0.18 (-2.84)** |
| Cruel to animals | **0.23 (6.54)** | **-0.14 (-2.45)** | **-0.15 (-2.44)** |
| Cruelty, bullying, or meanness to others | **0.45 (17.44)** | **-0.27 (-3.92)** | **-0.17 (-2.32)** |
| Daydreams or gets lost in his/her thoughts | **0.44 (21.74)** | **0.23 (3.72)** | **0.31 (4.45)** |
| Deliberately harms self or attempts suicide | **0.20 (6.10)** | 0.03 (0.43) | **-0.16 (-2.66)** |
| Demands a lot of attention | **0.58 (33.75)** | -0.01 (-0.11) | -0.06 (-0.79) |
| Destroys his/her own things | **0.53 (21.81)** | **-0.21 (-2.72)** | -0.09 (-1.18) |
| Destroys things belonging to his/her family or others | **0.56 (24.13)** | **-0.28 (-3.32)** | **-0.15 (-1.91)** |
| Disobedient at home | **0.57 (37.89)** | **-0.17 (-2.47)** | -0.04 (-0.56) |
| Disobedient at school | **0.51 (23.40)** | **-0.22 (-3.03)** | **0.14 (1.84)** |
| Doesn't eat well | **0.34 (15.47)** | 0.08 (1.56) | -0.03 (-0.71) |
| Doesn't get along with other kids | **0.50 (23.22)** | -0.12 (-1.53) | -0.00 (-0.05) |
| Doesn't seem to feel guilty after misbehaving | **0.50 (23.04)** | **-0.22 (-3.27)** | -0.05 (-0.67) |
| Easily jealous | **0.56 (32.86)** | -0.00 (-0.01) | -**0.17 (-2.37)** |
| Breaks rules at home, school or elsewhere | **0.60 (35.10)** | **-0.24 (-3.15)** | 0.00 (0.05) |
| Fears certain animals, situations, or places, other than school | **0.35 (15.11)** | **0.32 (6.74)** | **-0.15 (-2.99)** |
| Fears going to school | **0.31 (10.98)** | **0.29 (4.99)** | **-0.20 (-3.27)** |
| Fears he/she might think or do something bad | **0.33 (14.02)** | **0.37 (8.02)** | **-0.15 (-2.73)** |
| Feels he/she has to be perfect | **0.28 (12.98)** | **0.40 (11.09)** | **-0.25 (-5.55)** |
| Feels or complains that no one loves him/her | **0.50 (23.29)** | **0.15 (2.16)** | **-0.27 (-3.95)** |
| Feels others are out to get him/her | **0.41 (15.45)** | 0.11 (1.64) | **-0.23 (-3.48)** |
| Feels worthless or inferior | **0.49 (23.93)** | **0.32 (4.96)** | **-0.16 (-2.20)** |
| Gets hurt a lot, accident prone | **0.34 (13.93)** | **0.12 (2.31)** | -0.03 (-0.52) |
| Gets in many fights | **0.39 (12.59)** | **-0.26 (-3.33)** | **-0.19 (-3.00)** |
| Gets teased a lot | **0.45 (20.75)** | 0.08 (1.13) | 0.00 (0.02) |
| Hangs around with others who get in trouble | **0.38 (14.35)** | **-0.20 (-3.05)** | -0.01 (-0.11) |
| Hears sound or voices that aren't there | **0.23 (7.00)** | 0.07 (0.88) | **-0.10 (-2.26)** |
| Impulsive or acts without thinking | **0.63 (38.52)** | -0.08 (-1.02) | **0.16 (1.76)** |
| Would rather be alone than with others | **0.37 (17.28)** | **0.21 (3.74)** | -0.05 (-0.88) |
| Lying or cheating | **0.53 (28.75)** | **-0.19 (-2.74)** | 0.07 (0.90) |
| Bites fingernails | **0.24 (11.34)** | 0.06 (1.55) | **0.10 (2.27)** |
| Nervous, highstrung, or tense | **0.54 (29.70)** | **0.28 (3.77)** | **-0.16 (-2.28)** |
| Nervous movements or twitching | **0.41 (16.76)** | **0.16 (2.51)** | 0.04 (0.69) |
| Nightmares | **0.36 (16.01)** | **0.22 (3.97)** | **-0.13 (-2.40)** |
| Not liked by other kids | **0.48 (21.93)** | 0.02 (0.35) | 0.05 (0.72) |
| Constipated, doesn't move bowels | **0.28 (11.18)** | **0.19 (4.20)** | **-0.13 (-2.51)** |
| Too fearful or anxious | **0.45 (21.50)** | **0.47 (8.44)** | **-0.21 (-3.54)** |
| Feels dizzy or lightheaded | **0.26 (9.33)** | **0.24 (4.88)** | **-0.22 (-4.61)** |
| Feels too guilty | **0.32 (12.29)** | **0.42 (8.61)** | **-0.21 (-3.70)** |
| Overeating | **0.28 (10.69)** | **0.19 (3.60)** | -0.05 (-0.91) |
| Overtired without good reason | **0.33 (11.79)** | **0.22 (4.13)** | **-0.14 (-2.30)** |
| Overweight | **0.15 (6.12)** | **0.23 (6.43)** | -0.03 (-0.73) |
| Aches or pains (not stomach or headaches) | **0.25 (10.95)** | **0.24 (6.56)** | **-0.22 (-5.23)** |
| Headaches | **0.24 (11.50)** | **0.22 (6.03)** | **-0.25 (-7.12)** |
| Nausea, feels sick | **0.28 (11.24)** | **0.27 (6.20)** | **-0.27 (-6.60)** |
| Problems with eyes (not if corrected by glasses) | **0.16 (7.11)** | **0.13 (3.59)** | -0.07 (-1.64) |
| Rashes or other skin problems | **0.23 (10.20)** | **0.14 (3.43)** | **-0.17 (-4.33)** |
| Stomachaches | **0.31 (13.75)** | **0.28 (5.88)** | **-0.26 (-5.60)** |
| Vomiting, throwing up | **0.15 (5.78)** | **0.07 (1.80)** | **-0.12 (-3.40)** |
| Other (physical problems without known physical cause) | **0.15 (5.41)** | **0.13 (3.77)** | **-0.08 (-2.30)** |
| Physically attacks people | **0.43 (15.69)** | **-0.26 (-3.53)** | **-0.25 (-3.60)** |
| Picks nose, skin, or other parts of body | **0.38 (17.97)** | 0.04 (0.69) | 0.05 (0.84) |
| Plays with own sex parts in public | **0.16 (3.62)** | **-0.18 (-2.18)** | -0.05 (-0.79) |
| Plays with own sex parts too much | **0.19 (4.87)** | **-0.19 (-2.53)** | -0.06 (-0.88) |
| Poor school work | **0.47 (21.73)** | -0.04 (-0.64) | **0.23 (3.08)** |
| Poorly coordinated or clumsy | **0.45 (19.25)** | **0.18 (2.58)** | 0.08 (1.11) |
| Prefers being with older kids | **0.35 (15.30)** | 0.04 (0.79) | -0.01 (-0.22) |
| Prefers being with younger kids | **0.38 (17.62)** | **0.18 (3.45)** | **0.15 (2.31)** |
| Refuses to talk | **0.37 (15.67)** | **0.12 (1.87)** | **-0.09 (-1.67)** |
| Repeats certain acts over and over; compulsions | **0.44 (15.21)** | 0.06 (0.79) | 0.02 (0.29) |
| Runs away from home | **0.18 (4.37)** | **-0.20 (-2.36)** | **-0.18 (-3.31)** |
| Screams a lot | **0.56 (26.60)** | -0.12 (-1.40) | **-0.28 (-3.84)** |
| Secretive, keeps things to self | **0.43 (20.86)** | 0.09 (1.38) | **-0.12 (-1.94)** |
| Sees things that aren't there | **0.19 (6.29)** | 0.09 (1.35) | -0.07 (-1.53) |
| Self-conscious or easily embarrassed | **0.47 (27.77)** | **0.37 (6.46)** | -**0.20 (-3.35)** |
| Sets fires | **0.14 (4.83)** | **-0.16 (-3.27)** | -0.04 (-0.94) |
| Sexual problems | **0.18 (3.89)** | **-0.29 (-3.24)** | **-0.13 (-2.57)** |
| Showing off or clowning | **0.45 (22.98)** | -0.09 (-1.45) | 0.09 (1.34) |
| Too shy or timid | **0.30 (13.91)** | **0.32 (7.68)** | -**0.14 (-3.13)** |
| Sleeps less than most kids | **0.37 (14.17)** | **0.17 (2.62)** | -0.05 (-0.89) |
| Sleeps more than most kids during day and/or night | **0.23 (8.34)** | -0.00 (-0.01) | **-0.08 (-1.81)** |
| Inattentive or easily distracted | **0.63 (48.61)** | 0.11 (1.38) | **0.36 (3.88)** |
| Speech problem | **0.19 (7.52)** | 0.02 (0.40) | **0.07 (1.77)** |
| Stares blankly | **0.45 (18.37)** | **0.18 (2.85)** | **0.22 (2.89)** |
| Steals at home | **0.40 (13.48)** | **-0.29 (-4.14)** | -0.11 (-1.59) |
| Steals outside the home | **0.31 (9.16)** | **-0.31 (-4.71)** | -0.07 (-1.05) |
| Stores up too many things he/she doesn't need | **0.40 (17.69)** | 0.09 (1.53) | -0.02 (-0.37) |
| Strange behavior | **0.48 (18.13)** | -0.06 (-0.65) | -0.06 (-0.77) |
| Strange ideas | **0.44 (16.96)** | 0.10 (1.41) | 0.07 (0.97) |
| Stubborn, sullen, or irritable | **0.59 (37.08)** | 0.03 (0.36) | **-0.20 (-2.60)** |
| Sudden changes in mood or feelings | **0.61 (35.55)** | 0.06 (0.72) | **-0.25 (-3.00)** |
| Sulks a lot | **0.53 (24.84)** | **0.14 (1.91)** | **-0.23 (-3.08)** |
| Suspicious | **0.41 (14.13)** | -0.05 (-0.67) | -0.09 (-1.55) |
| Swearing or obscene language | **0.38 (13.43)** | **-0.16 (-2.49)** | **-0.16 (-2.67)** |
| Talks about killing self | **0.35 (11.01)** | **0.11 (1.70)** | **-0.27 (-4.29)** |
| Talks or walks in sleep | **0.20 (8.81)** | **0.09 (2.15)** | -0.05 (-1.18) |
| Talks too much | **0.43 (22.80)** | 0.06 (1.01) | **0.12 (1.80)** |
| Teases a lot | **0.43 (18.02)** | **-0.16 (-2.38)** | -0.08 (-1.20) |
| Temper tantrums or hot temper | **0.60 (37.73)** | -0.06 (-0.81) | **-0.26 (-3.30)** |
| Thinks about sex too much | **0.25 (6.02)** | **-0.24 (-2.56)** | **-0.22 (-3.68)** |
| Threatens people | **0.45 (14.30)** | **-0.29 (-3.18)** | **-0.34 (-4.72)** |
| Thumb-sucking | **0.08 (2.94)** | 0.03 (0.82) | -0.05 (-1.57) |
| Trouble sleeping | **0.44 (18.92)** | **0.25 (3.87)** | **-0.12 (-1.96)** |
| Truancy, skips school | **0.12 (3.54)** | 0.05 (0.82) | -0.06 (-1.61) |
| Underactive, slow moving, or lacks energy | **0.34 (11.97)** | **0.27 (4.69)** | -0.06 (-0.88) |
| Unhappy, sad, or depressed | **0.53 (24.01)** | **0.27 (3.87)** | **-0.29 (-3.96)** |
| Unusually loud | **0.51 (23.83)** | 0.02 (0.20) | -0.07 (-0.87) |
| Uses drugs for non medical purposes | 0.02 (0.66) | 0.00 (0.22) | -0.09 (-1.23) |
| Vandalism | **0.29 (8.01)** | **-0.19 (-2.69)** | -0.07 (-1.27) |
| Wets self during the day | **0.12 (3.97)** | -0.02 (-0.41) | 0.04 (0.90) |
| Wets the bed | **0.13 (5.43)** | -0.05 (-1.31) | -0.02 (-0.59) |
| Whining | **0.49 (26.15)** | 0.08 (1.30) | **-0.16 (-2.40)** |
| Wishes to be of opposite sex | **0.12 (3.60)** | 0.02 (0.20) | -0.02 (-0.58) |
| Withdrawn, doesn't get involved with others | **0.41 (16.16)** | **0.20 (3.17)** | -0.09 (-1.28) |
| Worries | **0.45 (25.36)** | **0.46 (8.67)** | **-0.25 (-4.28)** |

Supplementary file 1c. Pearson’s correlations between the structural and functional loadings. Significant correlations that survived FDR correction (*q* < 0.05) are indicated in **bold**.

| ***LC1*** | area | thickness | volume | between-network RSFC |
| --- | --- | --- | --- | --- |
| thickness | r = -0.04 p = 0.297 |  |  |  |
| volume | **r = 0.79 p = 0.000** | **r = 0.43 p = 0.000** |  |  |
| between-network RSFC | r = 0.14 p = 0.068 | r = -0.10 p = 0.147 | r = 0.15 p = 0.078 |  |
| within-network RSFC | **r = 0.11 p = 0.048** | **r = -0.13 p = 0.040** | r = -0.01 p = 0.460 | **r = -0.35**  **p = 0.000** |
| ***LC2*** | area | thickness | volume | between-network RSFC |
| thickness | r = 0.05 p = 0.275 |  |  |  |
| volume | **r = 0.83 p = 0.000** | **r = 0.53 p = 0.000** |  |  |
| between-network RSFC | r = 0.14 p = 0.105 | r = 0.16 p = 0.051 | **r = 0.24 p = 0.011** |  |
| within-network RSFC | r = -0.04 p = 0.333 | r = -0.08 p = 0.187 | r = -0.11 p = 0.089 | **r = -0.42**  **p = 0.000** |
| ***LC3*** | area | thickness | volume | between-network RSFC |
| thickness | r = -0.03 p = 0.399 |  |  |  |
| volume | **r = 0.92 p = 0.000** | **r = 0.32 p = 0.001** |  |  |
| between-network RSFC | r = 0.13 p = 0.106 | **r = 0.28 p = 0.004** | **r = 0.21 p = 0.019** |  |
| within-network RSFC | r = 0.07 p = 0.301 | r = -0.01 p = 0.512 | r = 0.07 p = 0.364 | **r = -0.45**  **p = 0.000** |

Supplementary file 1d. Pearson’s correlations between the principal gradient scores and PLS loadings by imaging modality. Significant correlations that survived FDR correction (*q* < 0.05) are indicated in bold. *Abbreviations: MID=Monetary Incentive Delay task; EN-BACK=Emotional N Back test; SST=Stop Signal task.*

| **LC** | **Modality** | **Pearson r** | **spin p** |
| --- | --- | --- | --- |
| LC1 | Area | 0.03 | 0.392 |
|  | Thickness | 0.09 | 0.182 |
|  | Volume | 0.00 | 0.531 |
|  | Within-network RSFC | **-0.14** | **0.010** |
|  | Between-network RSFC | **0.43** | **0.000** |
| LC2 | Area | 0.02 | 0.443 |
|  | Thickness | -0.01 | 0.421 |
|  | Volume | 0.01 | 0.482 |
|  | Within-network RSFC | **0.12** | **0.032** |
|  | Between-network RSFC | 0.11 | 0.060 |
| LC3 | Area | -0.03 | 0.429 |
|  | Thickness | -0.16 | 0.111 |
|  | Volume | -0.07 | 0.345 |
|  | Within-network RSFC | **0.17** | **0.017** |
|  | Between-network RSFC | **0.33** | **0.000** |

Supplementary file 1e. PLS loadings and z-scores for diffusion-based measures for LCs 1-3. Significant loadings and z-scores that survived FDR correction (*q* < 0.05) are indicated in bold.

| **DTI measure** | **LC1 loading (z-score)** | **LC2 loading (z-score)** | **LC3 loading (z-score)** |
| --- | --- | --- | --- |
| FA corpus callosum | 0.02 (0.57) | **0.06 (1.65)** | 0.06 (1.63) |
| FA foreceps major | 0.01 (0.40) | **0.06 (1.71)** | 0.02 (0.62) |
| FA foreceps minor | -0.01 (-0.32) | 0.06 (1.38) | **0.08 (1.76)** |
| FA left anterior thalamic radiations | **0.06 (1.70)** | -0.05 (-1.44) | **0.06 (1.75)** |
| FA left cingulate cingulum | -0.01 (-0.43) | -0.04 (-1.49) | **0.06 (2.09)** |
| FA left corticospinal/pyramidal | 0.04 (1.59) | **-0.06 (-2.20)** | **0.05 (1.79)** |
| FA left fornix | **0.06 (2.25)** | -0.04 (-1.41) | **0.06 (1.95)** |
| FA left inferior frontal superior frontal cortex | **0.05 (1.74)** | -0.01 (-0.28) | **0.06 (2.26)** |
| FA left inferior longitudinal fasciculus | **0.05 (1.88)** | 0.01 (0.41) | **0.06 (2.43)** |
| FA left inferior-fronto-occipital fasciculus | 0.03 (0.90) | -0.01 (-0.35) | 0.01 (0.47) |
| FA left parahippocampal cingulum | 0.03 (0.96) | -0.02 (-0.64) | **0.08 (2.50)** |
| FA left parietal superior longitudinal fasciculus | 0.01 (0.40) | -0.00 (-0.06) | **0.08 (2.82)** |
| FA left striatal inferior frontal cortex | -0.04 (-1.07) | 0.00 (0.11) | 0.03 (0.73) |
| FA left superior corticostriate | **0.09 (2.26)** | **-0.10 (-2.48)** | 0.04 (1.10) |
| FA left superior corticostriate-frontal cortex | **0.08 (2.01)** | **-0.11 (-2.63)** | 0.03 (0.81) |
| FA left superior corticostriate-parietal cortex | **0.09 (2.40)** | **-0.07 (-1.97)** | 0.05 (1.49) |
| FA left superior longitudinal fasciculus | 0.01 (0.24) | 0.03 (0.99) | **0.09 (2.94)** |
| FA left temporal superior longitudinal fasciculus | 0.01 (0.23) | 0.04 (1.42) | **0.08 (2.86)** |
| FA left uncinate | 0.00 (0.06) | -0.02 (-0.86) | **0.05 (1.97)** |
| FA right anterior thalamic radiations | 0.06 (1.62) | **-0.07 (-1.94)** | 0.04 (1.12) |
| FA right cingulate cingulum | -0.03 (-1.13) | -0.02 (-0.86) | **0.09 (3.19)** |
| FA right corticospinal/pyramidal | **0.08 (2.66)** | **-0.05 (-1.89)** | 0.02 (0.80) |
| FA right fornix | **0.09 (2.86)** | -0.02 (-0.64) | 0.03 (1.11) |
| FA right inferior frontal superior frontal cortex | **0.06 (1.91)** | -0.03 (-1.05) | **0.05 (1.74)** |
| FA right inferior longitudinal fasciculus | **0.07 (2.38)** | -0.03 (-1.16) | 0.04 (1.60) |
| FA right inferior-fronto-occipital fasciculus | 0.01 (0.28) | 0.00 (0.07) | -0.01 (-0.35) |
| FA right parahippocampal cingulum | 0.03 (1.00) | -0.02 (-0.75) | 0.02 (0.78) |
| FA right parietal superior longitudinal fasciculus | 0.02 (0.78) | 0.01 (0.28) | 0.02 (0.85) |
| FA right striatal inferior frontal cortex | -0.06 (-1.48) | 0.01 (0.14) | 0.02 (0.56) |
| FA right superior corticostriate | **0.12 (2.84)** | -0.08 (-1.77) | 0.01 (0.32) |
| FA right superior corticostriate-frontal cortex | **0.09 (2.34)** | **-0.09 (-2.15)** | -0.01 (-0.28) |
| FA right superior corticostriate-parietal cortex | **0.12 (3.17)** | -0.06 (-1.46) | 0.02 (0.63) |
| FA right superior longitudinal fasciculus | 0.02 (0.66) | 0.02 (0.77) | 0.02 (0.81) |
| FA right temporal superior longitudinal fasciculus | 0.01 (0.27) | **0.06 (2.18)** | 0.03 (0.96) |
| FA right uncinate | 0.00 (0.17) | 0.00 (0.06) | 0.03 (1.21) |
| MD corpus callosum | 0.07 (1.49) | -0.09 (-1.75) | -0.02 (-0.44) |
| MD foreceps major | 0.06 (1.24) | **-0.10 (-2.04)** | -0.04 (-0.76) |
| MD foreceps minor | **0.08 (1.69)** | -0.08 (-1.47) | -0.02 (-0.41) |
| MD left anterior thalamic radiations | 0.06 (1.22) | -0.07 (-1.50) | -0.02 (-0.42) |
| MD left cingulate cingulum | **0.08 (1.93)** | -0.05 (-1.24) | -0.02 (-0.54) |
| MD left corticospinal/pyramidal | 0.07 (1.37) | **-0.09 (-1.65)** | -0.03 (-0.58) |
| MD left fornix | 0.04 (0.85) | -0.08 (-1.62) | -0.04 (-0.70) |
| MD left inferior frontal superior frontal cortex | 0.04 (1.20) | -0.05 (-1.53) | -0.03 (-0.75) |
| MD left inferior longitudinal fasciculus | **0.08 (1.93)** | -0.06 (-1.45) | -0.06 (-1.32) |
| MD left inferior-fronto-occipital fasciculus | 0.07 (1.47) | -0.08 (-1.63) | -0.02 (-0.44) |
| MD left parahippocampal cingulum | **0.08 (1.72)** | -0.08 (-1.55) | -0.04 (-0.80) |
| MD left parietal superior longitudinal fasciculus | 0.03 (0.87) | -0.02 (-0.72) | -0.05 (-1.59) |
| MD left striatal inferior frontal cortex | **0.07 (1.74)** | -0.05 (-1.17) | -0.03 (-0.57) |
| MD left superior corticostriate | 0.06 (1.42) | -0.06 (-1.54) | -0.04 (-1.07) |
| MD left superior corticostriate-frontal cortex | 0.06 (1.55) | -0.06 (-1.50) | -0.03 (-0.83) |
| MD left superior corticostriate-parietal cortex | 0.05 (1.17) | -0.06 (-1.46) | -0.04 (-1.09) |
| MD left superior longitudinal fasciculus | 0.04 (1.24) | -0.06 (-1.59) | -0.06 (-1.61) |
| MD left temporal superior longitudinal fasciculus | 0.05 (1.37) | **-0.07 (-1.93)** | -0.06 (-1.55) |
| MD left uncinate | **0.08 (1.91)** | -0.06 (-1.30) | -0.02 (-0.53) |
| MD right anterior thalamic radiations | 0.06 (1.35) | -0.07 (-1.51) | -0.02 (-0.49) |
| MD right cingulate cingulum | **0.08 (1.84)** | -0.03 (-0.79) | -0.04 (-0.95) |
| MD right corticospinal/pyramidal | 0.06 (1.16) | **-0.09 (-1.80)** | -0.03 (-0.58) |
| MD right fornix | 0.03 (0.60) | **-0.09 (-1.72)** | -0.03 (-0.50) |
| MD right inferior frontal superior frontal cortex | 0.04 (1.30) | -0.04 (-1.08) | -0.02 (-0.54) |
| MD right inferior longitudinal fasciculus | **0.07 (1.76)** | -0.06 (-1.34) | -0.05 (-1.23) |
| MD right inferior-fronto-occipital fasciculus | 0.07 (1.48) | -0.08 (-1.56) | -0.03 (-0.61) |
| MD right parahippocampal cingulum | 0.07 (1.57) | **-0.09 (-1.96)** | -0.03 (-0.55) |
| MD right parietal superior longitudinal fasciculus | 0.02 (0.46) | -0.03 (-0.90) | -0.02 (-0.63) |
| MD right striatal inferior frontal cortex | 0.07 (1.56) | -0.06 (-1.38) | -0.04 (-0.82) |
| MD right superior corticostriate | 0.03 (0.77) | **-0.06 (-1.65)** | -0.03 (-0.84) |
| MD right superior corticostriate-frontal cortex | 0.05 (1.42) | -0.06 (-1.57) | -0.02 (-0.59) |
| MD right superior corticostriate-parietal cortex | 0.02 (0.65) | **-0.07 (-1.73)** | -0.03 (-0.76) |
| MD right superior longitudinal fasciculus | 0.02 (0.65) | -0.04 (-1.29) | -0.02 (-0.68) |
| MD right temporal superior longitudinal fasciculus | 0.04 (1.07) | **-0.08 (-2.08)** | -0.03 (-0.88) |
| MD right uncinate | 0.07 (1.61) | -0.07 (-1.51) | -0.02 (-0.46) |

Supplementary file 1f. Absolute Pearson’s correlations between original behavior/imaging loadings and loadings from control PLS analyses using PCA explaining 10%, 30%, 70%, 90% variance in each imaging modality. The number of principal components per imaging modality is shown for each control analysis. Significant correlations are indicated in bold. *Abbreviations: PCA=principal component analysis; PCs = principal components.*

|  |  | PCA explaining 10% variance | PCA explaining 30% variance | PCA explaining 70% variance | PCA explaining 90% variance |
| --- | --- | --- | --- | --- | --- |
|  | Surface area PCs | 4 | 21 | 102 | 226 |
|  | Thickness PCs | 1 | 11 | 144 | 281 |
|  | Volume PCs | 2 | 20 | 127 | 266 |
|  | RSFC PCs | 6 | 62 | 732 | 1900 |
|  | *Total PCs* | *13* | *114* | *1105* | *2673* |
| LC1 | Behavior loadings | **1.00** | **1.00** | **1.00** | **1.00** |
|  | Area loadings | **0.35** | **0.71** | **0.88** | **0.79** |
|  | Thickness loadings | **0.75** | **0.84** | **0.91** | **0.83** |
|  | Volume loadings | **0.61** | **0.80** | **0.89** | **0.80** |
|  | RSFC loadings | **0.58** | **0.88** | **0.92** | **0.86** |
| LC2 | Behavior loadings | **0.70** | **0.97** | **1.00** | **0.99** |
|  | Area loadings | **0.14** | **0.72** | **0.89** | **0.82** |
|  | Thickness loadings | **0.40** | **0.81** | **0.89** | **0.80** |
|  | Volume loadings | **0.19** | **0.78** | **0.89** | **0.81** |
|  | RSFC loadings | **0.38** | **0.76** | **0.87** | **0.78** |
| LC3 | Behavior loadings | **0.64** | **0.89** | **0.88** | **0.81** |
|  | Area loadings | **0.25** | **0.82** | **0.84** | **0.73** |
|  | Thickness loadings | 0.02 | **0.66** | **0.81** | **0.67** |
|  | Volume loadings | **0.35** | **0.81** | **0.84** | **0.73** |
|  | RSFC loadings | **0.03** | **0.78** | **0.82** | **0.68** |

Supplementary file 1g**.** Posthoc associations between age, age^2^ and sex, and subject-specific imaging or behavior scores for LCs 1-5. Associations were tested using Pearson’s correlations for continuous variables (age, age^2^) and using t-tests for categorical variables (sex). Tests that survived FDR correction (*q* < 0.05) are indicated in bold.

|  |  | Age | | Age^2^ | | Sex | | | |
| --- | --- | --- | --- | --- | --- | --- | --- | --- | --- |
|  |  | *r* | *p* | *r* | *p* | Mean (SD) in females | Mean (SD) in males | *t* | *p* |
| LC1 | Imaging scores | 0.00 | 0.936 | 0.00 | 0.950 | -0.25 (1.15) | 0.24 (1.22) | **-12.19** | **0.000** |
|  | Behavior scores | 0.02 | 0.304 | 0.02 | 0.293 | -0.47 (3.89) | 0.46 (4.67) | **-6.44** | **0.000** |
| LC2 | Imaging scores | **0.05** | **0.002** | **0.05** | **0.003** | 0.54 (1.18) | -0.53 (1.24) | **26.02** | **0.000** |
|  | Behavior scores | **0.04** | **0.024** | **0.04** | **0.028** | 0.40 (1.98) | -0.39 (1.92) | **12.02** | **0.000** |
| LC3 | Imaging scores | 0.00 | 0.933 | 0.00 | 0.905 | -0.31 (1.19) | 0.31 (1.27) | **-14.98** | **0.000** |
|  | Behavior scores | 0.02 | 0.214 | 0.02 | 0.233 | -0.16 (1.44) | 0.16 (1.79) | **-5.84** | **0.000** |

Supplementary file 1h. Child Behavior Checklist (CBCL) items used in PLS analysis.

| 1 | Acts too young for his/her age | 60 | Rashes or other skin problems |
| --- | --- | --- | --- |
| 2 | Drinks alcohol without parents' approval | 61 | Stomachaches |
| 3 | Argues a lot | 62 | Vomiting, throwing up |
| 4 | Fails to finish things he/she starts | 63 | Other (physical problems without known physical cause) |
| 5 | There is very little he/she enjoys | 64 | Physically attacks people |
| 6 | Bowel movements outside toilet | 65 | Picks nose, skin, or other parts of body |
| 7 | Bragging, boasting | 66 | Plays with own sex parts in public |
| 8 | Can't concentrate, can't pay attention for long | 67 | Plays with own sex parts too much |
| 9 | Can't get his/her mind off certain thoughts; obsessions | 68 | Poor school work |
| 10 | Can't sit still, restless, or hyperactive | 69 | Poorly coordinated or clumsy |
| 11 | Clings to adults or too dependent | 70 | Prefers being with older kids |
| 12 | Complains of loneliness | 71 | Prefers being with younger kids |
| 13 | Confused or seems to be in a fog | 72 | Refuses to talk |
| 14 | Cries a lot | 73 | Repeats certain acts over and over; compulsions |
| 15 | Cruel to animals | 74 | Runs away from home |
| 16 | Cruelty, bullying, or meanness to others | 75 | Screams a lot |
| 17 | Daydreams or gets lost in his/her thoughts | 76 | Secretive, keeps things to self |
| 18 | Deliberately harms self or attempts suicide | 77 | Sees things that aren't there |
| 19 | Demands a lot of attention | 78 | Self-conscious or easily embarrassed |
| 20 | Destroys his/her own things | 79 | Sets fires |
| 21 | Destroys things belonging to his/her family or others | 80 | Sexual problems |
| 22 | Disobedient at home | 81 | Showing off or clowning |
| 23 | Disobedient at school | 82 | Too shy or timid |
| 24 | Doesn't eat well | 83 | Sleeps less than most kids |
| 25 | Doesn't get along with other kids | 84 | Sleeps more than most kids during day and/or night |
| 26 | Doesn't seem to feel guilty after misbehaving | 85 | Inattentive or easily distracted |
| 27 | Easily jealous | 86 | Speech problem |
| 28 | Breaks rules at home, school or elsewhere | 87 | Stares blankly |
| 29 | Fears certain animals, situations, or places, other than school | 88 | Steals at home |
| 30 | Fears going to school | 89 | Steals outside the home |
| 31 | Fears he/she might think or do something bad | 90 | Stores up too many things he/she doesn't need |
| 32 | Feels he/she has to be perfect | 91 | Strange behavior |
| 33 | Feels or complains that no one loves him/her | 92 | Strange ideas |
| 34 | Feels others are out to get him/her | 93 | Stubborn, sullen, or irritable |
| 35 | Feels worthless or inferior | 94 | Sudden changes in mood or feelings |
| 36 | Gets hurt a lot, accident prone | 95 | Sulks a lot |
| 37 | Gets in many fights | 96 | Suspicious |
| 38 | Gets teased a lot | 97 | Swearing or obscene language |
| 39 | Hangs around with others who get in trouble | 98 | Talks about killing self |
| 40 | Hears sound or voices that aren't there | 99 | Talks or walks in sleep |
| 41 | Impulsive or acts without thinking | 100 | Talks too much |
| 42 | Would rather be alone than with others | 101 | Teases a lot |
| 43 | Lying or cheating | 102 | Temper tantrums or hot temper |
| 44 | Bites fingernails | 103 | Thinks about sex too much |
| 45 | Nervous, highstrung, or tense | 104 | Threatens people |
| 46 | Nervous movements or twitching | 105 | Thumb-sucking |
| 47 | Nightmares | 106 | Trouble sleeping |
| 48 | Not liked by other kids | 107 | Truancy, skips school |
| 49 | Constipated, doesn't move bowels | 108 | Underactive, slow moving, or lacks energy |
| 50 | Too fearful or anxious | 109 | Unhappy, sad, or depressed |
| 51 | Feels dizzy or lightheaded | 110 | Unusually loud |
| 52 | Feels too guilty | 111 | Uses drugs for non medical purposes (don't include alcohol or tobacco) |
| 53 | Overeating | 112 | Vandalism |
| 54 | Overtired without good reason | 113 | Wets self during the day |
| 55 | Overweight | 114 | Wets the bed |
| 56 | Aches or pains (not stomach or headaches) | 115 | Whining |
| 57 | Headaches | 116 | Wishes to be of opposite sex |
| 58 | Nausea, feels sick | 117 | Withdrawn, doesn't get involved with others |
| 59 | Problems with eyes (not if corrected by glasses) | 118 | Worries |
